# Speciation and gene flow across an elevational gradient in New Guinea kingfishers

**DOI:** 10.1101/589044

**Authors:** Ethan Linck, Benjamin G. Freeman, John P. Dumbacher

## Abstract

Closely related species with parapatric elevational ranges are ubiquitous in tropical mountains worldwide. The gradient speciation hypothesis proposes that these series are the result of *in situ* ecological speciation driven by divergent selection across elevation. Direct tests of this scenario have been hampered by the difficulty inferring the geographic arrangement of populations at the time of divergence. In cichlids, sticklebacks, and *Timema* stick insects, support for ecological speciation driven by other selective pressures has come from demonstrating parallel speciation, where divergence proceeds independently across replicated environmental gradients. Here, we take advantage of the unique geography of the island of New Guinea to test for parallel gradient speciation in replicated populations of *Syma* kingfishers that show extremely subtle differentiation across elevation and between historically isolated mountain ranges. We find that currently described high elevation and low elevation species have reciprocally monophyletic gene trees and form nuclear DNA clusters, rejecting this hypothesis. However, demographic modeling suggests selection has likely maintained species boundaries in the face of gene flow following secondary contact. We compile evidence from the published literature to show that while *in situ* gradient speciation in labile organisms such as birds appears rare, divergent selection and post-speciation gene flow may be an underappreciated force in the origin of elevational series and tropical beta diversity along mountain slopes.

## Introduction

Series of closely related species with parapatric elevational ranges are ubiquitous in tropical mountains worldwide, contributing to their globally high beta diversity (Diamond, 1972; Jankowski, Ciecka, Meyer, & Rabenold, 2009; Cadena et al., 2012; Terborgh & Weske, 1975). This striking biogeographic pattern is consistent with alternate hypotheses of how speciation proceeds in tropical forest faunas (Moritz, Patton, Schneider, & Smith, 2000). Under the gradient speciation hypothesis, elevational series are the result of local adaptation to divergent elevational niches that leads to the evolution of reproductive isolation *in situ* (i.e., ecological speciation without geographic isolation; Smith, Wayne, Girman, & Bruford, 1997; Moritz, Patton, Schneider, & Smith, 2000; Nosil, 2012; Caro, Caycedo-Rosales, Bowie, & Cadena, 2013; Beheregaray, Cooke, Chao, & Landguth, 2015). Alternatively, allopatric speciation followed by secondary contract and elevational range displacement through competition could lead to identical distributional patterns (Mayr, 1942; Endler, 1982; Cadena et al., 2012; Freeman, 2015).

Despite significant research attention, the origin of elevational series eludes easy synthesis. This is partly due to the difficulty of inferring the geographic mode of speciation and its evolutionary mechanisms from biogeographic and phylogenetic data alone (Endler, 1982; Losos & Glor, 2003). While the gradient speciation hypothesis predicts that lineages with parapatric elevational ranges will be sister to one another—a relationship found in *Ithioma* butterflies (Elias et al., 2009), *Leptopogon* flycatchers (Bates & Zink, 1994) and some Andean amphibians and reptiles (Arteaga et al., 2016; Guayasamin et al., 2017), but not in many other elevational series of Andean birds and mammals (e.g., Patton & Smith, 1992; Dingle, Lovette, Canaday, & Smith, 2006; Cadena et al., 2012; Caro, Caycedo-Rosales, Bowie, & Cadena, 2013; Cadena & Céspedes, 2019)—this pattern alone cannot distinguish divergence with geographic isolation from divergence without geographic isolation. Furthermore, both the lability of species ranges over evolutionary time and extinction can quickly obscure any phylogenetic signal of the geography of speciation (Losos & Glor, 2003). In contrast, population genomic data can provide information on rates of gene flow through time, and by proxy, the geographic mode of divergence (Moyle et al., 2017; Chapman, Hiscock, & Filatov, 2013; Chapman, Hiscock, & Filatov, 2016), but can only reveal the loci underpinning reproductive isolation (and by extension, its selective drivers) under particular circumstances (e.g., with exceptional sampling of hybrid zones).

To date, the strongest evidence for ecological speciation of any type has come from studies of parallel divergence on replicated selective gradients (Schluter & Nagle, 1995; Johannesson, 2002). The independent evolution of similar morphs of cichlids (Schliewen, Tautz, & Pääbo, 1994) and sticklebacks (Rundle, Nagel, Boughman, & Schluter, 2000) in response to similar ecological pressures convincingly demonstrates a link between local adaptation and the evolution of *in situ* reproductive isolation. In *Timema* stick insects, parallel adaptation to different host plants in allopatric populations provides a terrestrial analogue at small spatial scales (Nosil, Crespi, & Sandoval, 2002). While this approach evokes studies of phylogenetic relationships in clades with taxa that segregate by both latitude and elevation (e.g., Cadena & Céspedes, 2019), it differs in true geographic independence among replicates, putatively shallow timescales that reduce the probability that extinction and range shifts have obscured evolutionary signal, and phenotypic variance that is shared across similar environments. To our knowledge, only one study has explicitly tested predictions of parallel gradient speciation across elevation (Fuchs, Fjeldså, & Bowie, 2011).

Here, we take advantage of the unique geography of the island of New Guinea to perform such a test in replicated populations of interior forest kingfishers in the genus *Syma* that show extremely subtle differentiation across elevation and between geographic replicates. We collected genomic, morphological, and biacoustic data to evaluate species limits and evidence for assortative mating, phylogenetic relationships, and demographic history. Under a hypothesis of parallel gradient speciation, we predicted that highland and lowland populations of *Syma* in isolated mountain ranges that have never been connected by montane forest (Benz, 2011) would be sister to one another and more distantly related to their allopatric congeners. Alternately, under the secondary contact hypothesis, we predicted all highland and lowland populations would be reciprocally monophyletic. We additionally evaluated the role of gene flow during divergence in *Syma*, and compile evidence relevant to these hypotheses from the published literature.

## Materials and Methods

### Study system

As currently delimited, Yellow-billed and Mountain Kingfishers *Syma torotoro* and *S. megarhyncha* (Aves: Alcedinidae) are putative sister taxa that segregate by elevation (Pratt & Beehler, 2015; Diamond, 1972) across the island of New Guinea. Lowland forest species *S. torotoro* is reportedly smaller with a higher-pitched call, and primarily found below 700 m, while the slightly larger, deeper-voiced *S. megarhyncha* is primarily found above 1100 m to 2200 m or higher in mid-montane forest (Pratt & Beehler, 2015). A third population, Yellow-billed Kingfisher subspecies *S. t. ochracea*, may also merit species status; intermediate in size and divergent in call, it is restricted to the oceanic islands of the D’Entrecasteaux Archipelago in southeastern Papua New Guinea. (For the remainder of the manuscript we refer to this taxon as *S. (t.) ochracea* to reflect this uncertainty but will ignore subspecies level taxa in other cases.) All *Syma* taxa spend most of their time in the middle and upper strata of closed-canopy forests, where they feed on a mix of insects and small vertebrates (Pratt & Beehler, 2015).

While Mayr, Rand, and Diamond argued that elevational replacements in New Guinea form through secondary contact of differentiated lineages (Rand, 1936; Mayr, 1942; Diamond, 1972), some evidence suggests *Syma megarhyncha* and *S. torotoro* may instead have undergone gradient speciation across elevation. First, both mainland *Syma* are territorial and sedentary, making it plausible that dispersal was sufficiently reduced to be overcome by divergent selection across the steep environmental gradient they inhabit (Endler, 1977). Second, *S. megarhyncha’*s larger size (Beehler & Pratt, 2016) is consistent with predictions of morphological adaptation to a cooler climate (Freeman, 2017). Additionally, geographically isolated populations of *S. megarhyncha* in the mountains of the Huon Peninsula and the Central Ranges have fixed differences in bill markings, hinting they may have formed independently through parallel divergence from local *S. torotoro* populations. However, species limits and range-wide variation have never been quantitatively assessed with either phenotypic or genetic data and observed differences might instead reflect phenotypic plasticity or clinal variation of a single widespread lineage (Caro et al., 2013).

### Morphological and bioacoustics data

To evaluate phenotypic support for species limits, we measured bill length, bill depth, tarsus, wing chord, and tail length from 72 museum specimens of *Syma torotoro (n*=30), *S. (t.) ochracea* (*n*=10), and *S. megarhyncha* (*n*=32) at the American Museum of Natural History, collected from 1894-1965 and including only individuals of known sex as originally identified by the preparator. Using these data, we performed principal component analyses in R (R Core Team 2018) with log-transformed and normalized variables and used PC1 to build mixture models using the R package mclust v. 5.4.1, which we evaluated with a maximum likelihood classification approach (Scrucca, Fop, Murphy, & Rafferty, 2016). We downloaded all available vocalizations from *S. torotoro* (*n*=34) and *S. megarhyncha* (*n*=14) from xeno-canto and Cornell’s Macaulay Library. We filtered these data for quality and quantified 36 standard bioacoustic variables using the warbleR package v. 1.1.14 in R (Araya-Salas & Smith-Vidaurre, 2017), analyzing 278 distinct vocalizations from *S. torotoro* and 106 from *S. megarhyncha* in total. We ran PCA with normalized variables on the output and used these data to alternate species delimitation models using the same approach as with our morphological data.

### Sampling, library preparation, and DNA sequencing

To infer population genetic structure and phylogenetic history in *Syma*, we extracted DNA from fresh tissues (*n*=6) and toepad samples from historical museum specimens (*n*=34) from 28 individuals of *S. torotoro*, 2 individuals of *S. (t.) ochracea*, and 10 individuals of *S. megarhyncha* (*n*=10), including 3 individuals collected in 1928 by Ernst Mayr himself. Though only partially overlapping with samples used in morphometric analyses, these individuals represented the full extent of both species’ ranges in New Guinea and Australia (**Table S1**) and included all described subspecies. We extracted DNA from fresh tissues using a Qiagen DNAeasy kit and the manufacturer’s recommended protocol. For historical toepad samples (collected 1877-1973), we extracted DNA using either a using a phenol-chloroform and centrifugal dialysis method (Dumbacher & Fleischer, 2001) (for reduced representation sequencing) or a standard salt extraction protocol (for whole genome sequencing). Due to constraints of cost and time, we employed two complementary sequencing approaches. On a subset of samples (*n*=20), we performed reduced representation genome sequencing using a hybridization capture with RAD probes (hyRAD) approach, described in detail elsewhere (Linck, Hanna, Sellas, & Dumbacher, 2017). We sent the remaining samples to the UC Berkeley’s Vincent J. Coates Genomic Sequencing Laboratory, where laboratory staff prepared genomic libraries for low coverage whole genome sequencing (WGS) using Illumina TruSeq Nano kits and a modified protocol that enzymatically repaired fragments with RNAse and skipped sonication. They then pooled (*n*=20) and sequenced these samples with 150 base pair paired end reads on a single lane of an Illumina HiSeq 4000.

### Sequence assembly and variant calling

We processed demultiplexed reads from both sequencing strategies together with a custom bioinformatic pipeline optimized for handling degraded DNA data and available at https://github.com/elinck/syma_speciation. Briefly, we trimmed raw reads for adapters and low-quality bases using bbduk from BBTools version 38.06 suite of bioinformatics tools. We aligned these reads to an unpublished draft genome of Woodland Kingfisher *Halcyon senegalensis* from the Bird 10K Genome Project using bbmap with a k-mer value of 12, a maximum indel length of 200 bp, and a minimum sequence identity of 0.65. These highly sensitive alignment parameters were necessary given the clade containing Syma diverged from the clade containing *Halcyon* approximately 15 mya (Andersen, McCullough, Mauck, Smith, & Moyle, 2018). We used PicardTools v. 2.17.8 and GATK v. 3.6.0 (McKenna et al., 2010) to append read groups and perform local realignment on .bam files. We then used mapDamage 2.0.9 to account for postmortem damage to DNA from historical museum specimens by rescaling quality scores (Jónsson, Ginolhac, Schubert, Johnson, & Orlando, 2013). We performed multisample variant calling using the UnifiedGenotyper tool in GATK v. 3.6.0, and filtered our variant calls for missing data, coverage, and quality with VCFtools 0.1.16 (Danecek et al., 2011). We generated multiple SNP datasets by implementing different filters to suit the requirements and goals of different analyses; these settings and their underlying rationale are described in detail below. To complement our nuclear DNA sequences, we assembled near-complete and fully annotated mitochondrial genomes from a majority of individuals using mitofinder v. 1.0.2 (Allo et al., 2020) and a complete mtDNA genome from close relative *Todiramphus sanctus* as a reference (Andersen et al., 2015). We then extracted NADH dehydrogenase 2 (ND2) from these genomes and performed multiple sequence alignment using MAFFT v7.407 under its “–auto” parameter setting package (Katoh & Standley, 2013).

### Population structure inference

We evaluated population genetic structure within and between species using a nonparametric clustering approach. We performed principal component analysis of genotypes (PCA) and identified putative genetic clusters for *k*=2 through *k*=4 using adegenet v. 2.1.1 (Jombart, 2008) and a 95% complete dataset with 66,917 SNPs from 35 individuals of *Syma torotoro (n*=24), *S. (t.) ochracea* (*n*=2), and *S. megarhyncha* (*n*=9) that passed quality filters. This included individuals from both sequencing approaches with a minimum depth of coverage of 3x per individual, a maximum depth of coverage of 120x per individual, a minimum quality score of 30, and a minimum minor allele frequency of 0.05. These relatively lenient filtering parameters were chosen to permit the inclusion of samples from both sequencing strategies in our initial assessment of probable species limits. We evaluated the best-fit clustering model using the Bayesian Information Criterion, again implemented in adegenet. To ensure our results were not an artifact of different library preparation methods or tissue type, we employed two complementary approaches. First, we performed pairwise Wilcoxon rank sum tests evaluating whether differences in coverage or the proportion of missing data were significantly associated with inferred clusters. Second, we reran PCA using the same filters as above but instead 1) included only whole genome sequence data or 2) excluded all modern tissues. We then compared relationships among samples from these analyses with those observed in the full dataset.

### Phylogenetic inference

As we lacked whole genome sequences from an appropriate outgroup species, we evaluated phylogenetic relationships among using our mitochondrial DNA (ND2) alignment, which was near-complete and did not feature correlations between missing data and library preparation method or tissue type. We inferred a time-calibrated phylogeny in BEAST 2.6.0 (Bouckaert et al., 2019) from the 37 samples of *Syma torotoro (n*=24), *S. (t.) ochracea* (*n*=3), and *S. megarhyncha* (*n*=10) with sufficiently high quality ND2 sequence data, including Sacred Kingfisher *Todiramphus sanctus* as an outgroup (Andersen et al., 2015). We used a strict molecular clock with a rate of 2.9×10^-8^ following Lerner, Meyer, James, Hofreiter, & Fleischer (2011), a GTR+GAMMA model of nucleotide evolution, and a Yule species tree prior. We ran BEAST for 50 million generations, storing trees every 1000 generations, and assessed mixing, likelihood stationarity and adequate effective sample sizes (ESS) above 200 for all estimated parameters using Tracer v.1.7.1 (Rambaut, Drummond, Xie, Baele, & Suchard, 2018). We then generated a maximum clade credibility tree using TreeAnotator v.2.6.0, discarding the first 10% of trees as burnin (Bouckaert et al., 2019).

### Demographic inference

We inferred the demographic history of the diverging lineages identified in our clustering and phylogenetic analyses using moments v.1.0.0, which uses ordinary differential equations to model the evolution of allele frequencies (Jouganous, Long, Ragsdale, & Gravel, 2017). We used the joint_sfs_folded() function in Python package scikit-allel v.1.2.1 (Miles, Ralph, Rae, & Pisupati, 2019) to calculate a folded joint site frequency spectrum (JSFS) using a SNP dataset that included only whole genome sequencing data, a minimum minor allele count of 1, no more than 10% missing data per site, a minimum depth of 6x per individual, and a minimum quality score of 30. We chose these filtering parameters to reduce the influence of sequencing error without strongly biasing the JSFS. This dataset did not include *S. t. ochracea*, and had been further thinned to include only variants in approximate linkage equilibrium (defined as sites where *r^2^*<0.1) using scikit-allel’s locate_unlinked() function. To account for the remaining missing data, we projected the JSFS down to an effective sample size of 16 x 12 chromosomes for *S. megarhyncha* and *S. torotoro* respectively.

We then specified four basic demographic models chosen to represent plausible speciation scenarios in *Syma*, which differed by level and timing of gene flow. These were: an isolation-with-migration model (IM), which permitted gene flow throughout the history of diverging lineages; an isolation-with-ancestral-migration model (AM), which featured an initial period of gene flow followed by a period of isolation; a model of secondary contact (SC), which featured an initial period of isolation followed by a period of gene flow; and a model of strict isolation (SI), which allowed no gene flow following initial divergence. Next, we specified four additional models which differed from each of the above only by allowing exponential population growth in the most recent time period: an isolation-with-migration-and-growth model (IMg), an isolation-with-ancestral-migration-and-growth model (AMg), a secondary contact and growth model (SCg), and a strict isolation and growth model (SIg).

After initially optimizing parameters for fitting each model using the “optimize log” method and a maximum of 3 iterations, we ran 9 additional optimizations, randomly sampling starting values from a uniform distribution within the bounds of 1×10^-4^ and 10. We converted parameter values to real units using a genome-wide mutation rate of 2.3×10^-9^ (Smeds, Qvarnström, & Ellegren, 2016), an effective sequence length scaled to reflect our LD-thinned SNP dataset, and a generation time estimate of two years. We generated parameter uncertainty estimates for the best-fit demographic model (selected based on AIC values) by fitting it to 200 bootstrapped site frequency spectra generated using the fs.sample() function on our data, an approach which is appropriate when sites are unlinked (Jouganous et al., 2017).

### Literature review

Last, we reviewed the published literature for studies explicitly or implicitly testing predictions of the gradient speciation hypothesis in elevational series of birds. We first performed a Web of Science search on 29 May 2020 using the terms “speciation AND elevation* AND bird*”. From the 122 results, we retained 15 studies that met three critera: they (1) generated novel phylogenetic or population genetic data; (2) involved at least two congeneric species or otherwise classified reciprocally monophyletic lineages with different elevational ranges; and (3) focused on tropical or subtropical taxa. We augmented these results with 16 relevant studies missed by the initial search that we were aware of from other contexts or discovered through the cited literature. In sum, our review included studies addressing a total of 24 unique taxa (**Table 2**).

## Results and Discussion

Despite the aspects of the biology and distribution of *Syma* kingfishers that are consistent with parallel gradient speciation across elevation, phylogenetic and population genetic evidence lead us to unequivocally reject this hypothesis in *Syma*. A maximum likelihood phylogeny from the mitochondrial gene ND2 and a clustering analysis from genome-wide DNA sequence data indicate current species limits are largely correct: allopatric high-elevation populations of Mountain Kingfisher *Syma megahryncha* are indeed each other’s closest relative, as are all sampled populations of Yellow-billed Kingfisher *Syma torotoro* (**Figure 1b,d**). These results were robust to potential artifacts of sample type or sequencing strategy. PCA performed on only whole genome sequencing data or only historic samples recovered qualitatively similar patterns (**Figure S1**), and pairwise Wilcoxon rank sum tests found no statistically significant association between coverage or the proportion of missing data in samples (all *P*>0.05). Inferred nuclear DNA clusters and mtDNA lineages were also reflected by phenotypic data, occupying divergent though overlapping areas of principal component space (**Figure 1e,f**). The first principal component of all bioacoustic and morphological data was bimodally distributed with respect to *S. torotoro* and *S. megarhyncha*, as were individual trait measurements (**Figure S2a**), a pattern predicted under assortative mating (Cadena, Zapata, & Jiménez, 2018). Strong negative loadings on PC1 for all measured morphological variables are more consistent with general body size differences than divergent selection on a specific ecologically relevant trait (i.e., bill width) (**Table S3**).

**Figure 1.**
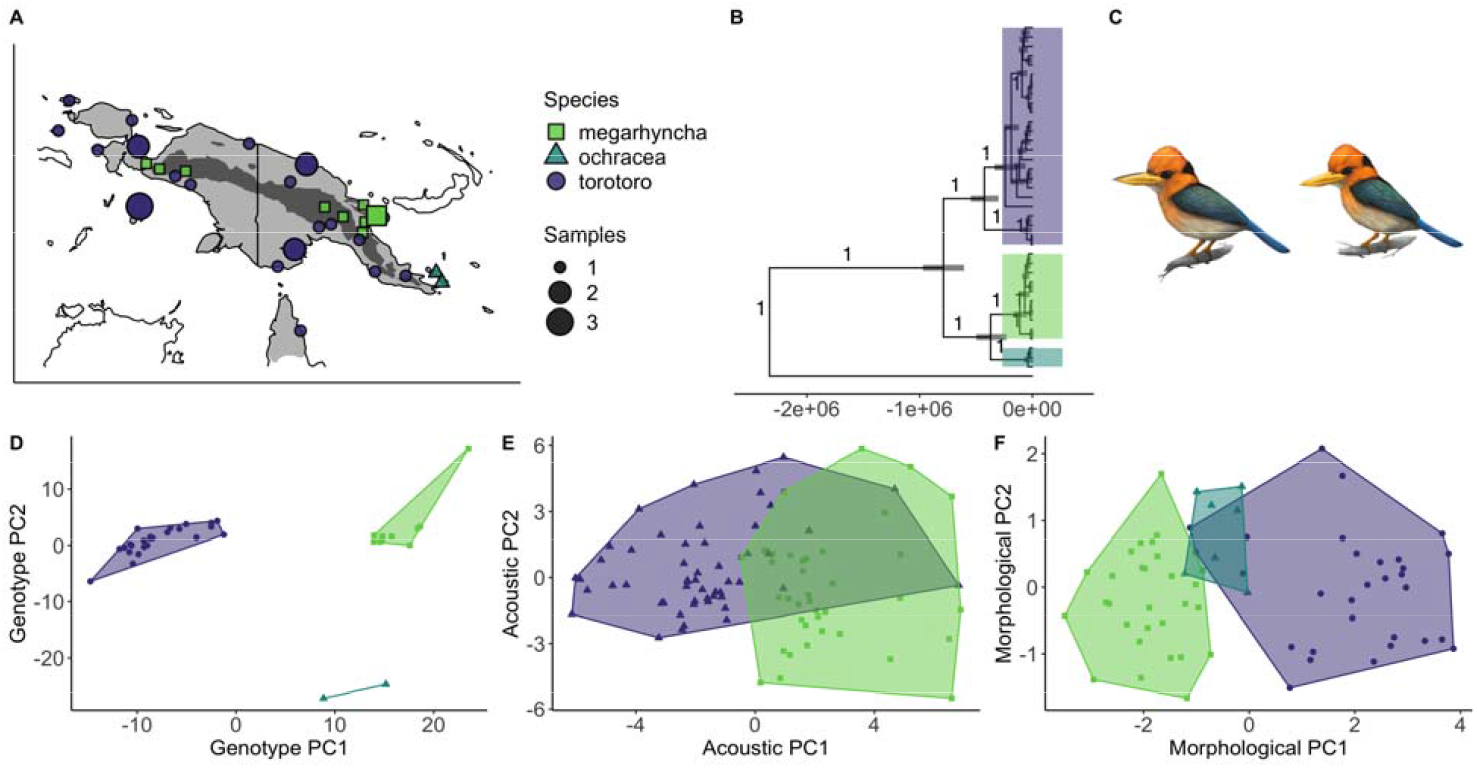
A) Sampling localities for *Syma* kingfishers across New Guinea and Australia, color-coded by genotype PC1 and scaled by number of individuals. B) A time-calibrated ND2 phylogeny supports reciprocal monophyly of *S. torotoro* and a clade with *S. megarhyncha* and *S. (t.)* ochracea, both themselves monophyletic. C) Illustration of *S. megarhyncha* (top) and *S. torotoro* (bottom), by Kevin Epperly. D). Principal component analysis of genotypes, clustered by the best fit *k*-means result (*K*=3) and color-coded by the mean PC1 value for all individuals **in** a given cluster. E) Principal component analysis of bioacoustic parameters, color-coded by species. F) Principal component analysis of morphological data, color-coded by species.

The clustering analysis and ND2 phylogeny alone do not exclude the possibility that gradient speciation occurred across a single slope followed by population expansion of the montane taxa to additional isolated montane regions. However, our best-fit demographic model of secondary contact (**Table 1**) is difficult to reconcile with speciation at the necessarily small spatial scale of a local mountainside. Furthermore, we note that our ND2 gene tree indicates an unexpected sister relationship between *S. megarhyncha* and the phenotypically distinctive Yellow-billed kingfisher subspecies *S. (t.) ochracea*, suggesting the latter may best be classified as a distinct biological species (**Figure 1b**). Though this relationship is consistent with multiple divergence histories—and reflects only the history of a single, nonrecombining locus—it presents a possible (if unparsimonious) scenario of allopatric speciation on an oceanic island followed by a subsequent reinvasion of the mainland and range displacement.

**Table 1.** Demographic model test results.

| Model | Log-likelihood | AIC |
| --- | --- | --- |
| Secondary contact (SC) | -564.062 | 1140.124 |
| Isolation-with-ancestral-migration (AM) | -590.0233 | 1192.047 |
| Strict isolation and growth (SIg) | -611.0493 | 1228.099 |
| Isolation-with-migration and growth (IMg) | -617.116 | 1244.232 |
| Isolation-with-migration (IM) | -631.8897 | 1273.779 |
| Strict isolation (SI) | -637.185 | 1280.37 |
| Isolation-with-ancestral-migration and growth (AMg) | -653.5472 | 1319.094 |
| Secondary contact and growth (SCg) | -716.0587 | 1444.117 |

In spite of the lack of evidence for parallel speciation, we emphasize our data nonetheless indicate a prominent role for natural selection in driving the evolution of elevational replacements. Parameter estimates from our model of secondary contact indicate *Syma torotoro* and *Syma megarhyncha* initially diverged in allopatry over 649,700 years ago (SD: 13,672 years), a value remarkably similar to the estimate from the time-calibrated ND2 gene tree (**Figure 1b**, 790,020 years ago; 95% CI: 607,421-972,618 years). This period of isolation was followed by a period of secondary contact and gene flow that initiated 250,000 years ago (**Table 1**). Our best estimate of the effective migration rate from *S. torotoro* to *S. megarhyncha* (*Nm*=1.14; SD: 0.02) is low but above the widely cited rule of thumb that one-migrant-per-generation prevents population stratification (Wright, 1931; Slatkin, 1985; Wang, 2004). This level of migration indirectly implies that some force—which we suggest is divergent selection— prevents gene flow from leading to the collapse of distinct populations (Nosil, 2012). Effective migration from *S. megarhyncha* to *S. torotoro* was considerably lower (*Nm*=0.11; SD=0.007), as expected given the lower inferred census population size of the smaller-ranged Mountain Kingfisher (*N*=368,390; SD=8,528) compared to the Yellow-billed Kingfisher (*N*=1,909,985; SD=26,028).

The apparent absence of recently admixed individuals in our dataset is unsurprising given the coarse grain of our sampling, but unfortunately precludes a rigorous assessment of the relative contribution of postzygotic (e.g., hybrid sterility) and prezygotic (habitat choice or assortative mating) isolation. However, the shallow divergence and long duration of gene flow between species leads us to suspect the former mechanism is at the very least only an incomplete reproductive barrier, suggesting extrinsic factors (particularly habitat requirements and selection against maladapted immigrants from different elevations) may be important. Evidence from recent field surveys that the species’ elevational range limits may not be truly parapatric in some cases but instead separated by hundreds of meters could further limit effective migration in either direction (Freeman & Freeman, 2014; Sam, Koane, & Novotny, 2014).

Ultimately, our assessment of speciation in *Syma* is consistent with trends revealed by a review of the literature on the origin of elevational series of tropical birds (**Table 2**). Of the 24 taxa included in our review, 17 concluded secondary contact was the exclusive mechanism behind the formational of elevational replacements. While seven taxa that showed patterns consistent with gradient speciation between at least two species, only a single study suggested gradient speciation was more common than allopatric divergence within elevational zones in a particular clade. All putative cases of gradient speciation lacked corroborating population genomic evidence. Of the eight taxa that had been explicitly tested for gene flow between elevational replacements, six detected it at appreciable levels. Thought this figure is likely inflated by ascertainment bias, it nonetheless suggests hybridization between elevational replacements may be more than previously appreciated (e.g., Cadena & Céspedes, 2019). We believe that while these findings confirm that divergence in allopatry likely generates the overwhelming majority of elevational series and support a shift away from *in situ* gradient speciation in labile organisms such as birds, they should reinforce a focus on divergent selection in the origin of tropical beta diversity. As the discovery of gene flow between diverging lineages at some point in the speciation continuum becomes the norm rather than the exception (Nosil, 2008; Kumar et al. 2017; Schumer et al., 2018; Edelman et al., 2019), we suggest that ecological and phenotypic differences that are evident during early divergence are likely to be adaptive and to function as drivers of the speciation process.

**Table 2.** Previous studies addressing the origin of elevational series of tropical birds using molecular data. The column “Speciation mode” describes the inferred model of divergence for each clade; when multiple species pairs in a study demonstrated conflicting histories, we describe the more common process but acknowledge heterogeneity using the word “mostly”. In some cases, we have assigned speciation mode based on our interpretation of the data rather than an explicitly stated conclusion by the original authors. The column “Gene flow” indicates whether gene flow between elevational replacements was tested for, and if so, whether or not it was detected.

| Citations | Taxon | Region | Speciation mode | Gene flow |
| --- | --- | --- | --- | --- |
| Bates & Zink, 1994 | *Leptogon*<br>flycatchers | Andes | Gradient<br>speciation | N/A |
| Arctander & Fjeldsa,<br>1994; Cadena et al.,<br>2020 | *Scytalopus*<br>tapaculos | Andes | Mostly secondary<br>contact | N/A |
| Garcia-Moreno et al.,<br>1998 | *Ochthoeca*<br>chat-tyrants | Andes | Secondary contact | N/A |
| Garcia-Moreno et al.,<br>1999a | *Cranioleuca*<br>spinetails | Andes | Secondary contact | N/A |
| Garcia-Moreno et al.,<br>1999b; Benham et<br>al., 2014 | *Metallura*<br>hummingbirds | Andes | Secondary contact | N/A |
| Garcia-Moreno et al.,<br>2001 | *Hemispingus*<br>tanagers | Andes | Secondary contact | N/A |
| Burns & Naoki, 2004 | *Tangara*<br>tanagers | Andes | Secondary contact | N/A |
| Dingle et al., 2006;<br>Caro et al., 2013;<br>Halfwerk et al.,<br>2016; Cadena et al.,<br>2019 | *Henicorhina*<br>wood wrens | Andes | Secondary contact | Yes |
| Ribas et al., 2007 | *Pionus* parrots | Andes | Secondary contact | N/A |
| Cadena, 2007 | *Buarremon* | Andes | Secondary contact | N/A |
|  | brush-finches |  |  |  |
| Chaves et al., 2007 | *Adelomyia melanogenys* hummingbird lineages | Andes | Possibly gradient speciation | Yes |
| Norman et al., 2007 | *Meliphaga* honeyeaters | Australasia | Mostly secondary contact | N/A |
| Parra et al., 2009 | *Coeligena* hummingbirds | Andes | Secondary contact | N/A |
| Sedano & Burns, 2010 | 93 tanager species | Andes | Mostly secondary contact | N/A |
| Fuchs et al., 2011 | *Phyllastrephus debilis* lineages | Afrotropics | Secondary contact | Yes |
| Päckert et al., 2011 | Seven passerine groups | Himalaya | Secondary contact | N/A |
| Dubay & Witt, 2012; Dubay & Witt, 2014 | *Anareites* tit-tyrants | Andes | Secondary contact | Yes |
| Wu et al., 2014 | Leiotherichinae babblers | Himalaya | Secondary contact | N/A |
| Winger et al., 2014 | *Grallaria antpittas* | Andes | Secondary contact | N/A |
| Voelker et al., 2015 | *Pheonicurus* redstarts | Himalaya | Mostly secondary contact | N/A |
| Moyle et al., 2017 | Five pairs of passerine taxa | Borneo | Secondary contact | No |
| Morales-Rozo et al., 2017 | *Ramphocelus* tanagers | Andes | Secondary contact | Yes |
| Cowles & Uy, 2019 | *Zosterops* white-eyes | Australasia | Secondary contact | No |
| Gabrielli et al., 2020; Bourgeois et al., 2020 | *Zosterops bourbonicus* white-eyes | Reunion Island | Possibly gradient speciation | Yes |

## Data availability

All code used in this study can be found at https://github.com/elinck/syma_speciation. Processed data are available from Dryad [doi: 10.5061/dryad.4f4qrfj9b], and sequence data are available from the NCBI SRA [accession: pending].

## Acknowledgements

We thank generations of New Guinean landowners and field assistants without whose help this paper would have been impossible to write. For tissue loans, we thank L. Joseph at ANWC, K. Zykowski at YPM, R. Moyle and M. Robbins at KUMNH, P. Sweet at AMNH, and R. Prys-Jones at NHMUK-Tring. For help arranging fieldwork, we thank A. Mack, G. Kapui, J. Robbins, N. Gowep, B. Beehler, L. Dabek, N. Whitmore, F. Dem, S. Tulai, B. Iova., and V. Novotny. We thank A. Wiebe for assistance measuring specimens and thank C.J. Battey for his many contributions. This work was supported by NSF Doctoral Dissertation Improvement Grant #1701224 to J. Klicka and E.B.L, a NDSEG Fellowship to E.B.L., and by NSF DEB #0108247 to J.P.D

## Supplemental material

**Table S1.** Sampling, sequencing, and analysis information. WP=Indonesian New Guinea; PNG=Papua New Guinea.

| Specimen | Species | Subspecies | Locality | Tissue source | Date collected | Sequencing strategy | PCA | mtDNA phylogeny | Demographic inference |
| --- | --- | --- | --- | --- | --- | --- | --- | --- | --- |
| AMNH:Birds:293714 | torotoro | torotoro | Ifaar, WP | toepad | 1928 | hyRAD | yes | yes | no |
| AMNH:Birds:293715 | torotoro | torotoro | Kepaur, WP | toepad | 1897 | hyRAD | yes | yes | no |
| AMNH:Birds:300723 | torotoro | torotoro | Kepaur, WP | toepad | 1897 | hyRAD | yes | yes | no |
| AMNH:Birds:300723 | torotoro | torotoro | Misol Island, WP | toepad | 1900 | hyRAD | yes | yes | no |
| AMNH:Birds:329542 | torotoro | torotoro | Wasior, WP | toepad | 1928 | hyRAD | yes | yes | no |
| AMNH:Birds:437798 | torotoro | torotoro | Amberbaki, WP | toepad | 1877 | hyRAD | yes | yes | no |
| AMNH:Birds:637429 | torotoro | torotoro | Humbolt Bay, WP | toepad | 1928 | hyRAD | yes | yes | no |
| AMNH:Birds:637441 | torotoro | torotoro | Mt. Mori, WP | toepad | 1899 | hyRAD | yes | yes | no |
| AMNH:Birds:637445 | torotoro | torotoro | East Sepik Province, PNG | toepad | 2003 | hyRAD | yes | yes | no |
| AMNH:Birds:637446 | torotoro | torotoro | Waigeu Island, WP | toepad | 1900 | hyRAD | yes | yes | no |
| AMNH:Birds:637450 | torotoro | tentelare | Aru Islands, WP | toepad | 1896 | hyRAD | yes | yes | no |
| AMNH:Birds:637464 | torotoro | tentelare | Aru Islands, WP | toepad | 1900 | hyRAD | yes | yes | no |
| AMNH:Birds:637464 | torotoro | ochracea | Normanby Island, PNG | toepad | 1934 | hyRAD | yes | yes | no |
| CAS:Birds:7131 | torotoro | pseuestes | Gulf Province, PNG | muscle | 2002 | hyRAD | yes | yes | no |
| KU:Birds:5215 | torotoro | pseuestes | Gulf Province, PNG | muscle | 2003 | hyRAD | yes | yes | no |
| KU:Birds:5464 | torotoro | pseuestes | Gulf Province, PNG | muscle | 2003 | hyRAD | yes | yes | no |
| KU:Birds:626 | torotoro | meeki | Varirata National Park, PNG | muscle | 2011 | hyRAD | yes | yes | no |
| KU:Birds:6927 | torotoro | meeki | Mt. Suckling, PNG | muscle | 2011 | hyRAD | yes | yes | no |
| NHMUK:Birds:1911.12.20.822 | torotoro | pseustes | Satakwa River, WP | toepad | 1911 | hyRAD | yes | yes | no |
| NHMUK:Birds:1911.12.20.823 | torotoro | pseustes | Mimika River, WP | toepad | 1913 | hyRAD | yes | yes | no |
| AMNH:Birds:301861 | megarhyncha | wellsi | Weylendgep Kunupi, WP | toepad | 1931 | WGS | yes | yes | yes |
| AMNH:Birds:302859 | megarhyncha | wellsi | Mt. Derimapa, WP | toepad | 1930 | WGS | yes | yes | yes |
| AMNH:Birds:339957 | megarhyncha | wellsi | Bernhard Camp, WP | toepad | 1939 | WGS | yes | yes | yes |
| AMNH:Birds:808986 | megarhyncha | megarhyncha | Okapa, PNG | toepad | 1965 | WGS | yes | yes | yes |
| ANWC:Birds:B02192 | megarhyncha | megrhyncha | Western Highlands, PNG | toepad | 1963 | WGS | yes | yes | yes |
| ANWC:Birds:B04293 | megarhyncha | megarhyncha | Morobe, PNG | toepad | 1966 | WGS | yes | yes | yes |
| ANWC:Birds:B25307 | megarhyncha | megarhyncha | Wagau, PNG | toepad | 1973 | WGS | yes | yes | yes |
| ANWC:Birds:B25652 | megarhyncha | sellamontis | Huon Peninsula, PNG | toepad | 1973 | WGS | yes | yes | yes |
| ANWC:Birds:B26140 | megarhyncha | sellamontis | Huon Peninsula, PNG | toepad | 1973 | WGS | yes | yes | yes |
| YPM:Birds:91444 | megarhyncha | sellamontis | Huon Peninsula, PNG | toepad | 1969 | WGS | yes | yes | yes |
| AMNH:Birds:329539 | torotoro | ochracea | Fergusson Island, PNG | toepad | 1928 | WGS | yes | yes | yes |
| AMNH:Birds:329540 | torotoro | ochracea | Fergusson Island, PNG | toepad | 1928 | WGS | yes | yes | yes |
| AMNH:Birds:426095 | torotoro | flavirostris | Fly River, PNG | toepad | 1936 | WGS | yes | yes | yes |
| AMNH:Birds:426096 | torotoro | flavirostris | Fly River, PNG | toepad | 1936 | WGS | yes | yes | yes |
| AMNH:Birds:426121 | torotoro | meeki | Wassi Kussi River, PNG | toepad | 1937 | WGS | yes | yes | yes |
| AMNH:Birds:637460 | torotoro | tentelare | Aru Islands, WP | toepad | 1900 | WGS | yes | yes | yes |
| AMNH:Birds:637471 | torotoro | meeki | Simbang, PNG | toepad | 1899 | WGS | yes | yes | yes |
| AMNH:Birds:637511 | torotoro | flavirostris | Cape York Peninsula, AU | toepad | 1913 | WGS | yes | yes | yes |
| ANWC:Birds:B07293 | torotoro | torotoro | East Sepik Province, PNG | toepad | 1966 | WGS | yes | yes | yes |
| ANWC:Birds:B07499 | torotoro | torotoro | East Sepik Province, PNG | toepad | 1966 | WGS | yes | yes | yes |

**Table S2.** The proportion of missing data (genotypes) and mean fold sequencing coverage by library preparation method and tissue type. Missing data rates calculated from the 95% complete SNP matrix used in *k*-means clustering analyses.

| Sample subset | Missing data (±SD) | Sequencing coverage (±SD) |
| --- | --- | --- |
| WGS libraries | 0.007 (±0.010) | 6.022x (±3.567x) |
| hyRAD libraries | 0.015 (±0.013) | 4.427x (±2.795x) |
| Modern tissues | 0.007 (±0.009) | 5.452x (±3.321x) |
| Toepad tissues | 0.011 (±0.013) | 4.427x (±3.235x) |

**Table S3.** PCA loadings of log-transformed morphological variables for all *Syma* taxa.

| Trait | PC1 | PC2 | PC3 |
| --- | --- | --- | --- |
| Bill (from nostril) | -0.4249207 | 0.33505714 | 0.22963905 |
| Bill width | -0.4367190 | 0.27088119 | 0.08777821 |
| Bill depth | -0.4389784 | 0.14310077 | 0.30742375 |
| Tarsus | -0.3046142 | -0.85803722 | 0.37960606 |
| Wing chord | -0.4467254 | 0.05852709 | -0.16954328 |
| Tail length | -0.3790295 | -0.23287113 | -0.81988160 |

**Figure S1.**
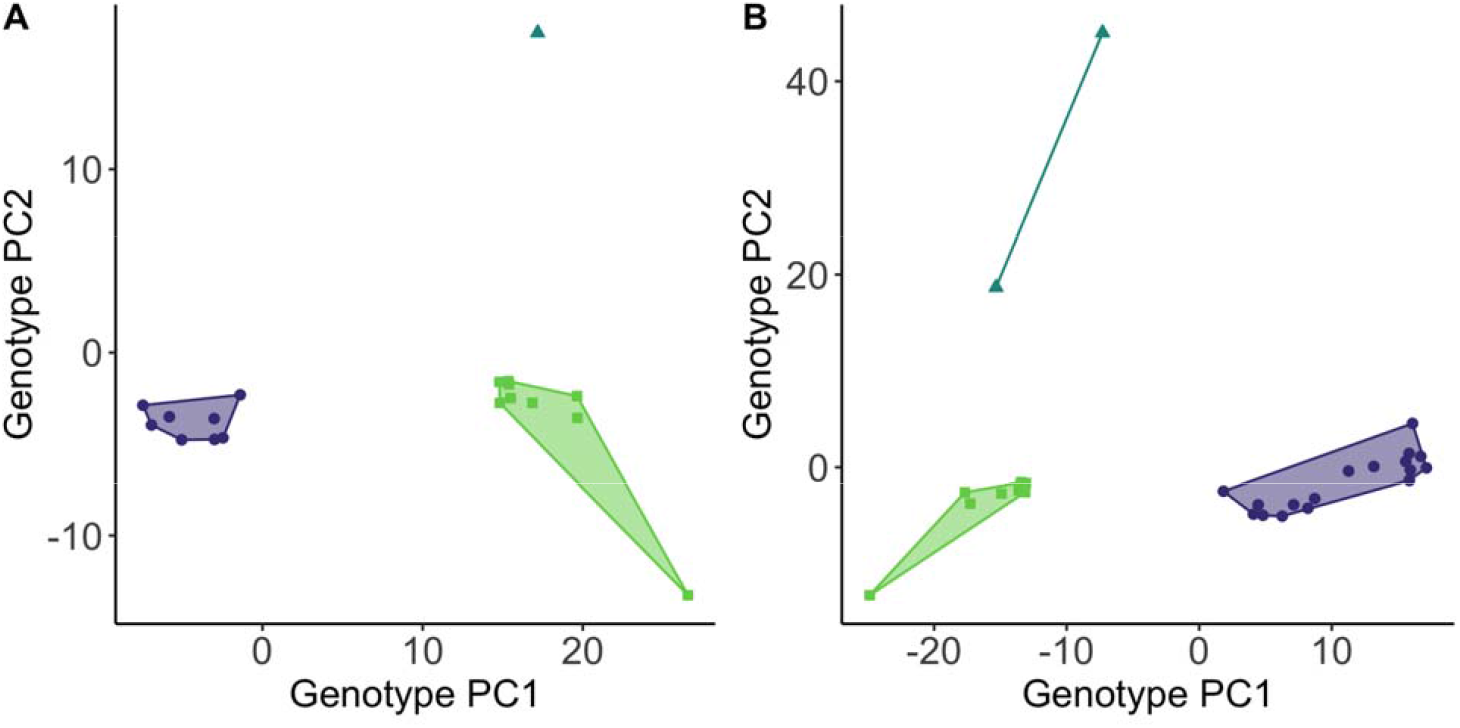
Principal component analysis of genotypes from A) whole-genome sequence data alone or B) only historic samples regardless of sequencing method, color-coded to match *k*-means results in Figure 1D. Note the single *S. (t) ochracea* individual high on PC2 in panel A.

**Figure S2.**
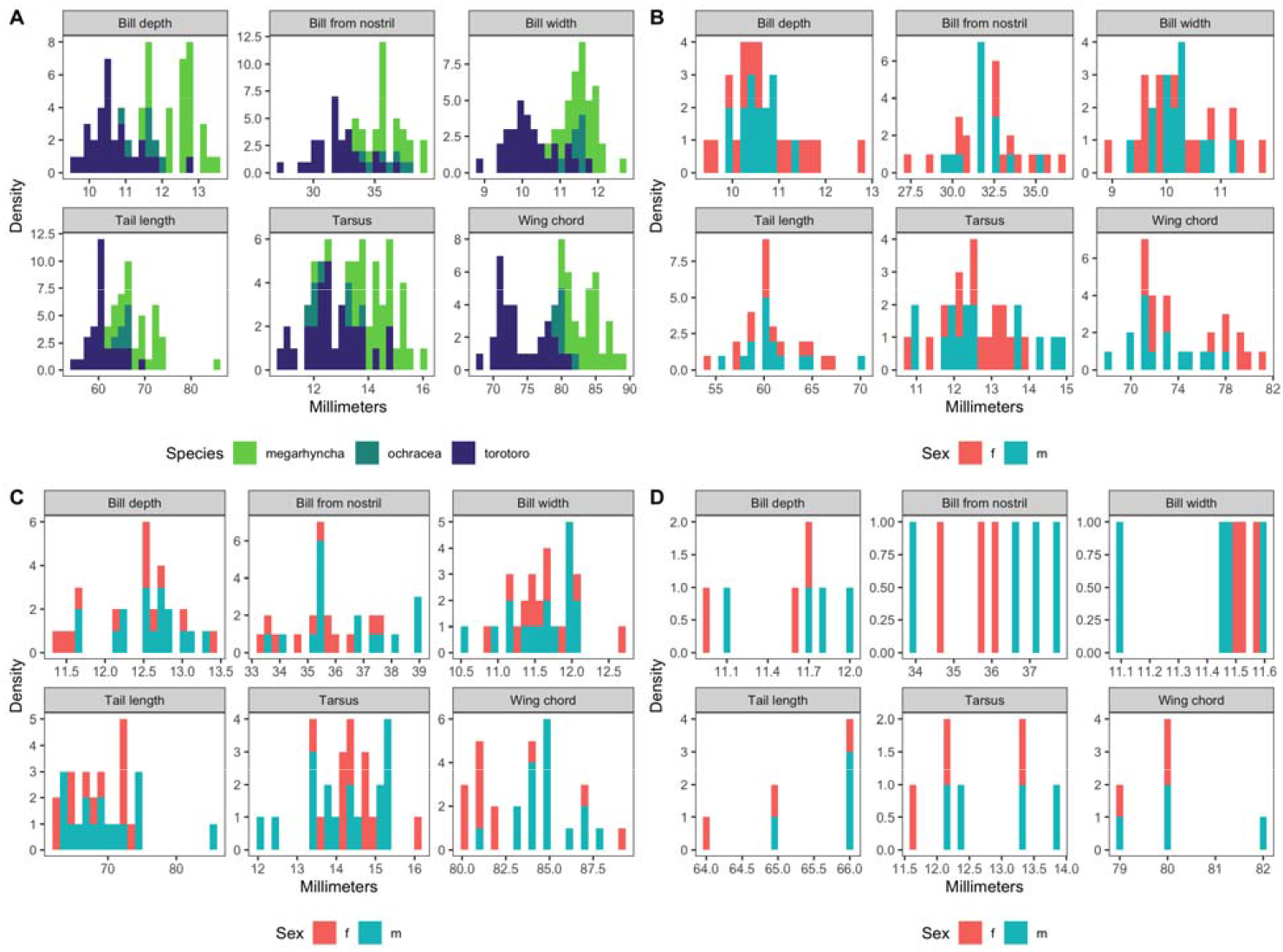
A) The distribution of log-transformed morphological trait measurements by taxon. B-D) The distribution of log-transformed morphological trait measureements by sex for *S. tororoto, S. megarhyncha*, and *S. (t.) ochracea*, respectively. Note the limited sample size of *S.(t.) ochracea*.

